# Increased microglial synapse elimination in patient-specific models of schizophrenia

**DOI:** 10.1101/231290

**Authors:** Carl M. Sellgren, Jessica Gracias, Bradley Watmuff, Carleton P. Goold, Jessica M. Thanos, Ting Fu, Rakesh Karmacharya, Hannah E. Brown, Jennifer Wang, Steven D. Sheridan, Roy H. Perlis

## Abstract

Schizophrenia patients display decreased synaptic density in postmortem studies, suggesting aberrant microglial synapse elimination during neurodevelopment. Here, we use cellular reprogramming to create patient-specific *in vitro* models of microglia-mediated synapse engulfment that demonstrate increased synapse elimination in schizophrenia-derived models compared to healthy controls. We show that excessive synaptic pruning in schizophrenia reflects abnormalities in microglia-like cells as well as synaptic structures. Further, we find that schizophrenia risk-associated variants within the complement component 4 locus contribute to the increased uptake in schizophrenia models. Finally, we demonstrate that the antibiotic minocycline reduces microglia-mediated synapse uptake and show that minocycline treatment for acne is associated with a reduction in incident schizophrenia risk compared to other treatments in a cohort of more than 9,000 young adults drawn from health records. Specific pharmacological interventions targeting excessive pruning merit further study for their capacity to delay or prevent the onset of schizophrenia in high-risk individuals.

## INTRODUCTION

Schizophrenia is a heritable psychiatric disorder with disabling psychotic and cognitive symptoms ^1^. Disease mechanisms remains largely elusive despite the identification of more than 100 regions of the genome associated with schizophrenia liability ^2^, repeated imaging studies displaying reduced grey matter thickness as well as abnormal functional connectivity ^3–11^, and postmortem studies reporting reduced numbers of synaptic structures ^12–14^. The most strongly associated locus for schizophrenia in meta-analysis of genome-wide association data involves structurally distinct alleles of the complement component 4 (*C4*) genes ^2^, with each allele associated with schizophrenia risk in proportion to its effect on *C4A* expression ^15^, and rodent studies suggest a pivotal role for complement signaling in microglia-mediated elimination of synapses in the developing visual system ^15–17^. Given the extensive elimination of synapses in human cerebral cortex during late adolescence and early adulthood ^18^, i.e., the time-period in which schizophrenia symptoms usually become clinically evident ^1^, it has been suggested that excessive synaptic pruning by microglia contributes to the observed reduction in synapse density among schizophrenia patients ^15,19^. However, the inability to model microglial functions in human using patient-derived cells has limited efforts to directly link excessive microglial synapse elimination to schizophrenia pathology. Mouse models cannot fully address this question, as the rodent genome does not code specialized *C4A* and *C4B* genes, and human microglia differ markedly from their rodent counterparts, with a higher expression of complement components as well as several genes important in brain development ^20^.

We have recently developed and validated high-throughput methods for modeling synaptic pruning *in vitro* using patient-derived cells ^21^. Similar to recent induced pluripotent stem cell (iPSC)-derived microglia-like cells ^22–25^, human monocytes cultured under conditions resembling the central nervous system (CNS) microenvironment recapitulate important traits of primary human microglia including a ramified morphology, expression of genes highly enriched in acutely isolated microglia (e.g., *P2ry12, C1qa, Gas6, Mertk, Gpr34*, and *Pros1*), and clustering with primary fetal microglia rather than peripheral monocyte-derived macrophages ^21^. Mimicking primary fetal microglia, induced microglia-like cells (iMGs) under serum-free *in vitro* conditions engulf synapses isolated from induced iPSC-derived neural cultures using a complement 3 receptor (C3R)-dependent mechanism ^21^. By using live-imaging, as well as confocal microscopy, we have been able to develop robust methods for quantification of synapse uptake ^21^.

In this study, we show that co-culture of iMGs with differentiated human neurons results in decreased spine density, and that iMGs display an increased uptake of isolated synaptic structures from schizophrenia-derived neurons compared to those from healthy controls (HCs). Further, increased synapse elimination among schizophrenia patients can be attributed to both the target synaptic structures as well as iMG function, and these effects are modified by, but not fully accounted for, variation within *C4A* and *C4B* present in the neural cultures. Finally, we demonstrate that minocycline, a second-generation tetracycline with high brain penetrance, reduces synapse uptake *in vitro* in a dose-responsive fashion. In electronic health records we likewise detect the predicted reduction in incident schizophrenia risk associated with minocycline exposure among acne-treated adolescents compared to other antibiotics.

## RESULTS

### Patient-derived microglia-like cells

To derive iMGs from schizophrenia subjects and matched HCs, monocytes were isolated from blood drawn from male individuals with schizophrenia and age-matched male HCs. Briefly, microglial induction was achieved by exposure to interleukin (IL)-34 and granulocyte macrophage colony stimulating factor (GM-CSF), under serum-free culture conditions, on an extracellular matrix (ECM) resembling astrocyte-derived ECM, containing laminin, collagen and entactin ^21^. No major differences in morphology could be observed between patient and control derived cultures with the vast majority of cells displaying a ramified morphology resembling resting-state microglia (**Fig. 1a**), and expressing microglia markers such as P2ry12, TMEM119, and PU.1 (**Fig. 1b**). To further characterize iMGs we performed transcriptome analyses comparing mRNA expression levels in iMGs to matched monocytes (**Fig. 1c and Supplementary Fig. 1**), as well as monocyte-derived macrophages (GM-CSF, 10 *%* fetal bovine serum [FBS]; **Fig. 1d**, and **Supplementary Fig. 2**). In comparison to monocytes (*P* < 0.0001), as well as to ordinary monocyte-derived macrophages (*P* < 0.0001), iMGs displayed a strong enrichment of microglia-specific genes (based on two gene sets of published microglia-specific genes in acutely isolated microglia ^20,26^; **Supplementary Tables 1, 2, 3** and **4**) among upregulated genes (fold change > 20), with iMGs significantly clustering apart from monocytes and monocyte-derived macrophages (**Fig. 1e** and **Supplementary Fig. 3** [displaying a hierarchical cluster analysis with uncertainty in clustering assessed by multiscale bootstrap resampling]). Corroborating our previous observations with isolated synapses from HCs ^21^, we also noted a clear decrease in spine density after co-culturing iMGs with iPSC-derived neural cultures (**Fig. 1f** and **1g**) while total numbers of neurons were unchanged.

**Figure 1.**
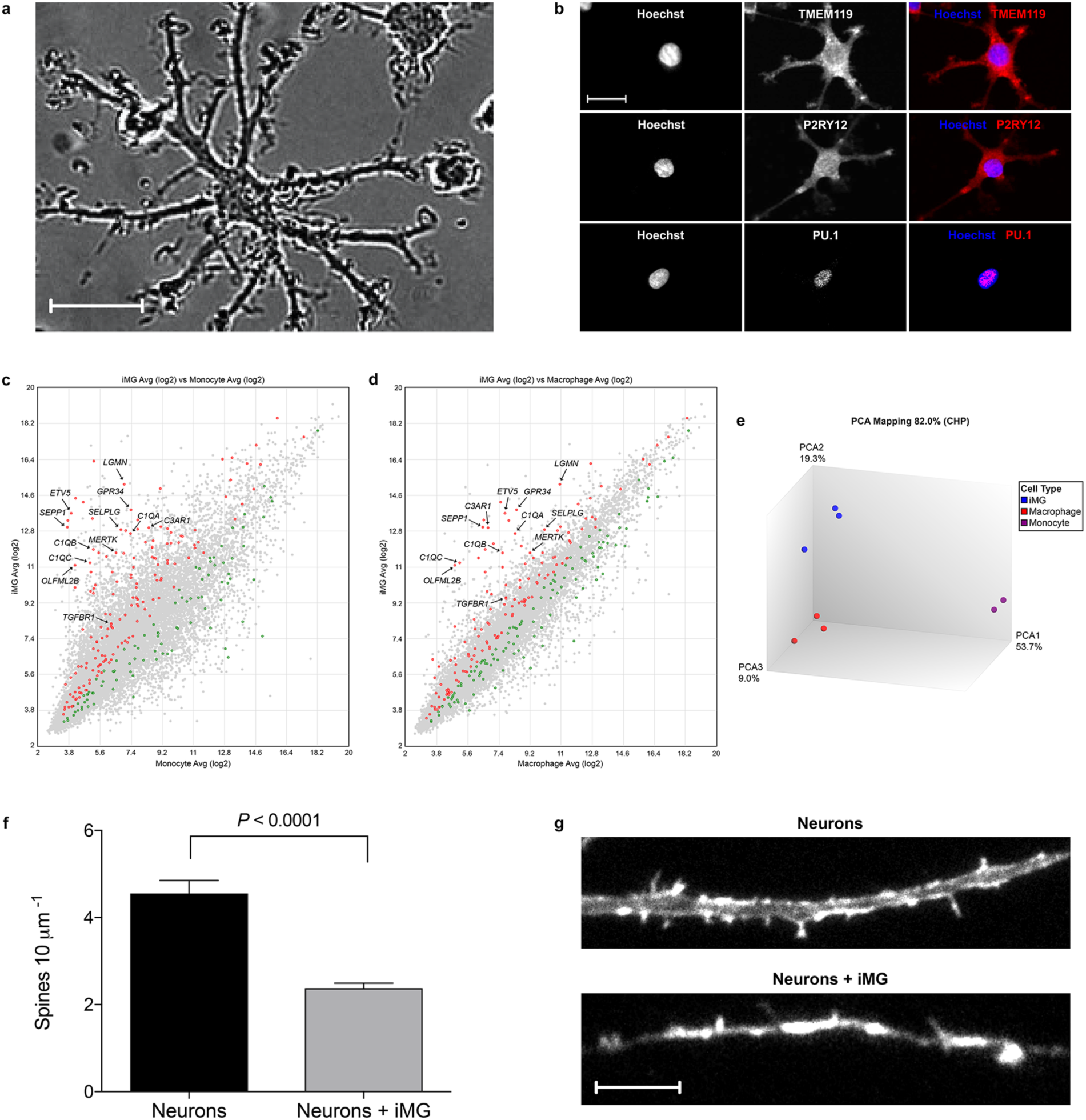
Characterizations of induced microglia-like cells (iMG). (**a**) Representative phase-contrast image of iMGs captured during a live-imaging session. Scale bar 20μm. (**b**) Confocal images of iMGs stained for TMEM119, P2RY12, and PU.1. Scale bar 20μm. (**c**) Scatter plot depicting mRNA expression levels in human monocytes (n=2) and matched iMGs (n=3). Red (higher expression in iMG) and green (higher expression in monocytes) indicate microglia-specific genes as reported by Bennett et al. ^26^. (**d**) Scatter plot depicting mRNA expression levels in human monocyte-derived macrophages (GM-CSF, 10 % fetal bovine serum [FBS], n=3) and matched iMG (n=3). Red (higher expression in iMGs) and green (higher expression in monocyte-derived macrophages) indicate microglia-specific genes as reported by Bennett et al. ^26^. (**e**) PCA of monocytes, monocyte-derived macrophages, and iMG cell populations based on mRNA expression levels. See also Fig. S3 for hierarchical cluster analysis using the R package pvclust. (f) Spine counts in an iPSC-derived neural line (Male, 37 years old) before and after cocultured with iMGs (n=7) for 48 hours. (g) Representative images of Phalloidin 647-stained dendrite of an iPSC-derived neuron before and after being co-cultured with iMGs for 24 h. Scale bar 10 μm.

### Isolation of active synapses from human iPSC-derived neural cultures

To model interactions between synapses and microglia in schizophrenia, we collected and reprogrammed iPSCs (**Fig. 2a**, and **Supplementary Fig. 4**) from dermal fibroblasts from male subjects with schizophrenia and age-matched HC donors, with clinical status confirmed by structured clinical interview, using an integration-free and all-synthetic RNA-based reprogramming system as previously described^27^. Purified and expandable neural progenitor cells (NPCs) were derived from these iPSCs using a neural induction supplement based on defined small molecule induction ^28^, followed by magnetic-activated cell sorting (PSA-NCAM^+^, CD271^−^, CD133^+^) and verified by immunocytochemical analyses for standard NPC markers (**Fig. 2b**). We then performed an inclusive neural differentiation by mitogen removal resulting in mixed populations of neural subtypes including telencephalic cortical (**Fig. 2c, 2d**, and **Supplementary Table 5**) ^29,30^. To be able to model interactions between iMGs and synapses in patients vs. HCs, we then proceeded with isolation of synaptosomes, i.e., synaptic nerve terminals. As we have previously shown, iPSC-derived neural cultures yields a highly purified synaptosome preparation compared to isolation from postmortem human brain ^21^. Briefly, neural cultures were harvested, homogenized and purified using Syn-PER Synaptic Protein Extraction Reagent (Thermo Scientific), which was followed by repeated centrifugal fractionation. In addition to previous validation of iPSC-derived synaptosomes using western blotting analyses for enrichment of synaptic markers ^21^, we now confirmed that synaptosomes after freeze-thaw exhibited canonical structural and functional synaptic properties by using transmission electron microscopy (TEM; **Supplementary Fig. 5**), labeling for active synaptic vesicle endocytosis (SVE) upon depolarization (**Fig. 2e**) ^31^, and fixation followed by staining for the post-synaptic marker PSD-95 (**Fig. 2e**). These investigations confirmed specificity of the synaptosome extraction procedure as well as functionality of isolated and freeze-thawed synaptosomes, and further enabled control for line-specific differences of synaptosome yield in functional assays.

**Figure 2.**
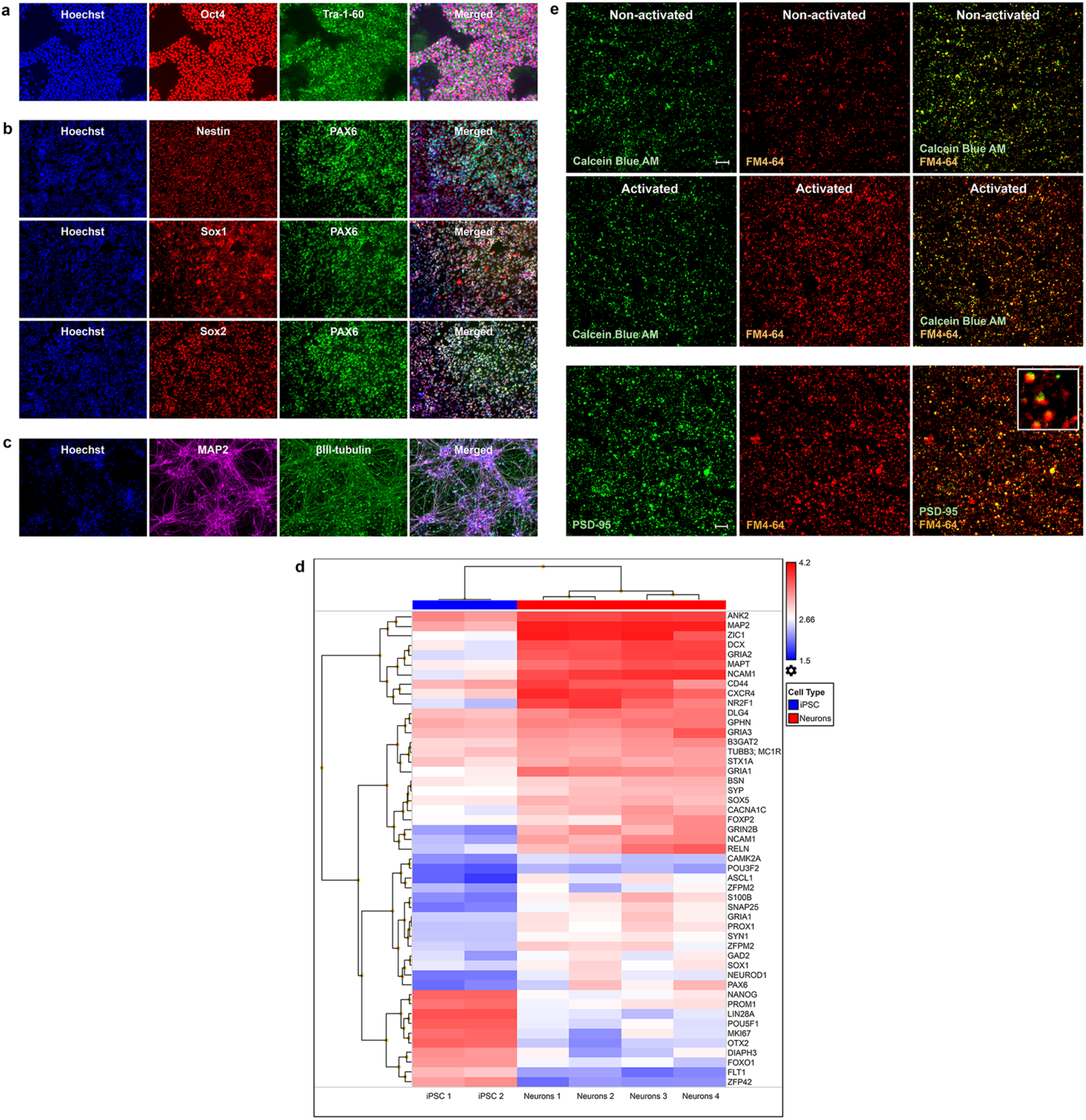
Isolation of active synaptic structures from iPSC-derived neural cultures. Representative confocal images of (**a**) induced pluripotent stem cells (iPSCs), (**b**) iPSC-derived neural progenitors (NPCs) and (**c**) neurons stained for indicated identity markers. (**d**) Heat map of mRNA expression values determined in iPSCs (n=2) and iPSC-derived neurons (n=4); for details see Table S5. (**e**) Confocal images of synaptosomes labeled with calcein blue–AM and FM4-64 (first and second row). Synaptosomes were activated by the addition of 40 mM KCl as indicated. Last row display synaptosomes labeled with FM4-64 and the synaptic marker PSD-95. Scale bar 60 μm.

### Increased engulfment of synapses in disease models – effect of potential confounding variables and *C4-HERV* copy numbers

We first compared overall phagocytosis in models incorporating iMGs and synaptosomes derived from 3 males with schizophrenia to models derived from 2 age-matched male HCs in real-time live imaging using synaptosomes labeled with a pH-sensitive dye (pHrodo^®^) that binds non-specifically to proteins and exclusively fluoresces bright red in the post-phagocytic phagolysosome compartments (**Fig. 3a**). Each synaptosome line was assessed with iMG cultures from at least 4 different subjects (in total, iMGs from 13 schizophrenia patients and 13 HCs) matched on age. Quantification of bright red fluorescence, indicating cellular uptake of pHrodo-labeled synaptosomes, revealed significantly increased phagocytosis in schizophrenia-derived iMG cultures compared to matched HC iMG cultures (*P* < 0.0001; **Fig. 3b**). As previously described ^21^, the vast majority of bright red fluorescence also stained positive for the post-synaptic marker PSD-95 (**Fig. 3c**) confirming that we detected a specific engulfment of synaptosomes. Using confocal microscopy we quantified phagocytic inclusions (PSD-95^+^ inclusions, 0.5–1.5 μm) in these cultures (excluding 2 iMG culture in which immunohistochemistry failed because of technical issues), as well as another set of iMG cultures consisting of 8 schizophrenia models and 6 HC models (iMG derived from 2 subjects with schizophrenia and 4 HCs, and synaptosomes derived from 2 schizophrenia subjects and 2 HCs). In agreement with real-time live imaging data, experiments quantifying PSD-95^+^ phagocytic inclusions using confocal microscopy confirmed an increased uptake in schizophrenia-derived models (*P* = 0.021; **Fig. 3d**).

**Figure 3.**
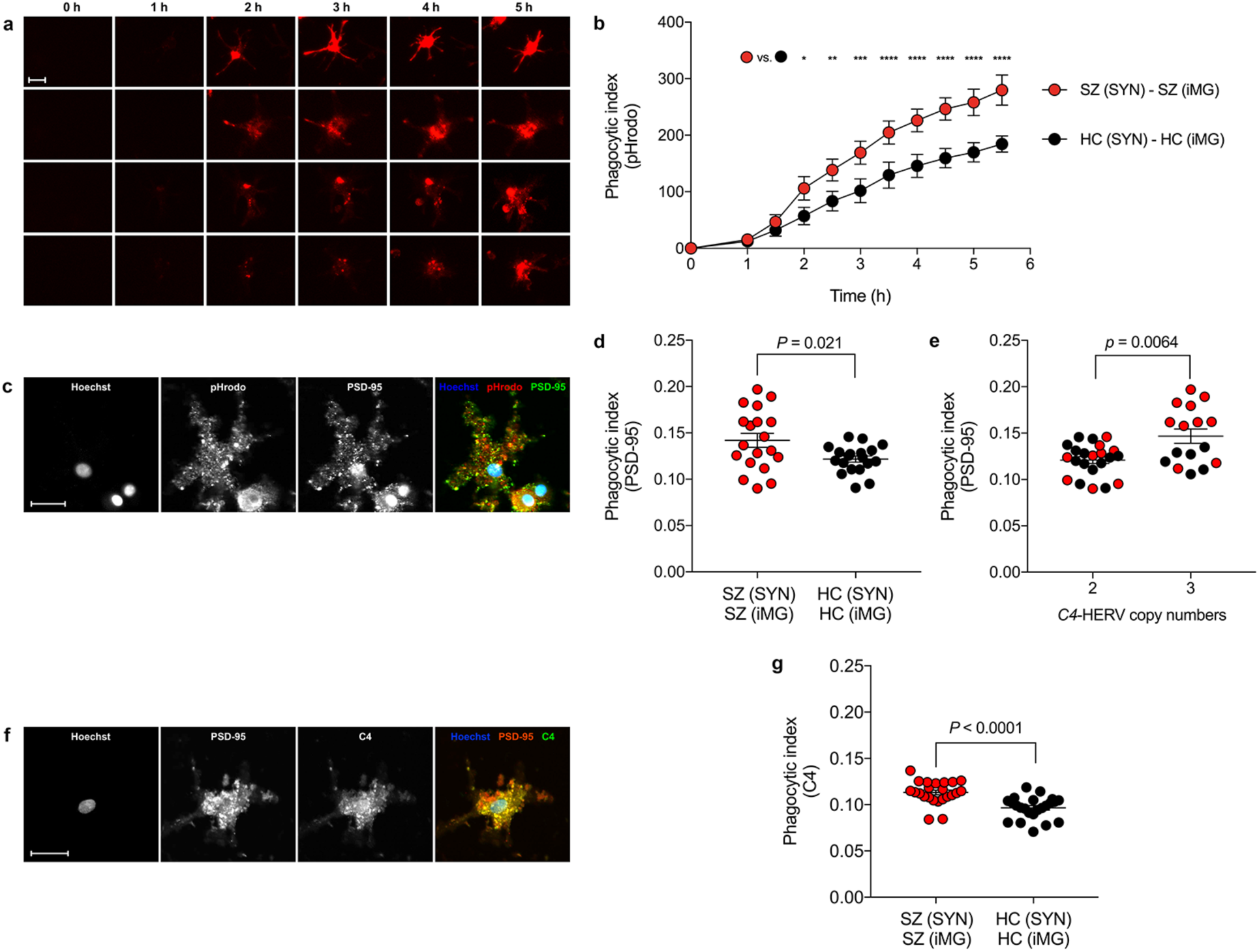
Increased engulfment of synaptic structures in schizophrenia-derived models. (**a**) Synaptosomes (SYN) were labeled with a pH-sensitive dye (pHrodo^®^) that fluoresces bright red in the post-phagocytic phagolysosome compartments and uptake was measured as red fluorescence. Induced microglia-like cell (iMG) cultures mixed with labeled SYN were imaged and red fluorescence was measured over 5.5 hours in real time using an IncuCyte ZOOM system (Essen Bioscience). Nine images (20x) per well were taken every 30 min. For more details regarding quantification see Online Methods. (**b**) Quantification of pHrodo (red)-labeled SYN uptake in iMGs during live imaging sessions of 5.5 h. Red represents mean for schizophrenia (SZ)-derived models (iMGs derived from 13 subjects and SYN derived from 3 subjects) and black for matched healthy control (HC; iMGs derived from 13 subjects and SYN derived from 2 subjects) models. Phagocytic index represents pHrodo+ area per iMG cell. (**c**) Confocal images showing that the vast majority of pHrodo^+^ puncta co-localize with the post-synaptic marker PSD-95. Scale bar 20μm. (**d**) Quantification of phagocytic inclusions (PSD-95^+^ inclusions, 0.5–1μm) using confocal microscopy and in an expanded sample containing SYN derived from a total of 5 patients and 4 HCs, matched with iMGs from 13 patients and 17 HCs (19 SZ models and 19 HC models). (**e**) Quantification of PSD-95^+^ inclusions for the same subjects but divided on C*4*-HERV copy numbers. Red denotes patients and black HCs. (**f**) SYN engulfed by iMGs, here indicated as PSD-95^+^ inclusions, strongly co-localized with C4^+^ staining. Costes’ P value for colocalization = 1. Scale bar 20 μm. (**g**) Quantification of phagocytic inclusions (C4^+^ inclusions, 0.5-1.5 μm) in 23 SZ models (SYN derived from 5 patients and iMGs derived from 11 patients) and 23 HC models (SYN derived from 5 HCs and iMGs derived from 18 HCs) using confocal microscopy All reported p-values are two-sided and error bars represent SEM. Data in (c) were analyzed using a two-way repeated repeated measures ANOVA with Sidaks’s post tests, and data in (d), (e), and (g) were analyzed using i-tests not assuming equal SDs.

To address potential clinical confounding variables, we first examined IQ as it was significantly lower in the schizophrenia group (see also **Supplementary Table 6**). We identified no association between iMG uptake of pHrodo-labeled synaptosomes and total IQ across all subjects (*r* [Pearson] = -0.27 *P* = 0.18). Likewise, iMG uptake of pHrodo-labeled synaptosomes was neither associated with clozapine treatment (54% of the schizophrenia group, OR = 1.01 [95% CI: 0.99-1.03]), nor use of an antidepressant (46% of the schizophrenia group, OR = 0.98 [95% CI: 0.94-1.02]).

If structural variation within *C4A* and *C4B* increases risk of schizophrenia via more extensive tagging of synapses by C4A, in turn leading to increased pruning, we reasoned that an effect of such genetic variation would influence overall synapse engulfment in our *in vitro* assay. We therefore used digital droplet PCR (ddPCR) to determine total copy numbers of *C4A* and *C4B* with human endogenous retroviral (HERV) insertion (*C4*-HERV [*C4L*]); shown to accurately predict schizophrenia risk originating from this locus as well as *C4A* over *C4B* expression ^15^. As previously described, synaptosomes prepared from astrocyte-conditioned medium (ACM) treated iPSC-derived neural cultures contains C4 protein (**Supplementary Fig. 6**). Stratifying PSD-95^+^ uptake in the total of 19 schizophrenia models and 19 HC models dependent on C4-HERV copy numbers (in neural lines used to isolate synaptosomes) we observed a higher uptake with increasing *C4*-HERV copy numbers (**Fig. 3e**) independent of total *C4* copy numbers (β = 0.025 [*P* = 0.0023] and adjusted β = 0.038 [*P* = 0.00037], respectively). We also observed an interaction between C4-HERV copy numbers and case status predicting higher PSD-95^+^ uptake (β = 0.043 [*P* = 0.0034]). The effect remained after adjusting for *C4*-HERV copy numbers and total *C4* copy numbers in matched iMGs (β = 0.042 [*P* = 0.022] while C4 haplotypes of iMGs had no impact on PSD-95^+^ uptake (*C4*-HERV; *P* = 0.99 and *C4* total; *P* = 0.87, respectively).

Confirming our prior observations^21^, engulfed synaptosomes (indicated as PSD-95^+^ inclusions) also strongly co-localized with C4^+^ puncta (Costes’ P value for colocalization = 1; **Fig. 3f**), thereby further supporting that engulfed synaptosomes were largely tagged by C4A. In line with this observation, schizophrenia models displayed increased uptake of C4^+^ synaptosomes (*P*<0.0001; **Fig. 3g**).

### Microglial factors influence synapse engulfment independent of neural phenotype

In order to better dissect the individual contributions of synaptosomes and iMGs toward the observed increase in phagocytic uptake of patient-derived synaptic particles, we repeated phagocytosis experiments but now *crossing* schizophrenia-derived synaptosomes with iMGs from a subset of individuals with schizophrenia as well as with synaptosomes from matched HCs. Specifically, we compared synapse elimination in “pure” disease models, i.e., iMG cultures from 8 subjects with schizophrenia matched with synaptosomes from 2 subjects with schizophrenia, with “crossed” or “mixed” models in which the same synaptic structures from schizophrenia subjects were added to iMGs derived from 8 matched HCs, as well as to matched “pure” control models (synaptosomes derived from 3 HCs; **Fig. 4a**). These experiments revealed a distinct and significantly decreased uptake of diseased synaptosomes when switching from schizophrenia-to HC-derived iMG (real-time live imaging: *P* < 0.0001; **Fig. 4b**, and confocal microscopy [PSD-95+ puncta] as well as C4^+^ puncta: *P* = 0.031; **Fig. 4c** and **Fig. 4d**), while in agreement with concurrent neural specific effects an increased uptake remained in mixed models compared to pure HC models. That is, both neural and microglial factors contribute to the overall increased uptake of synaptic structures in our schizophrenia models.

**Figure 4.**
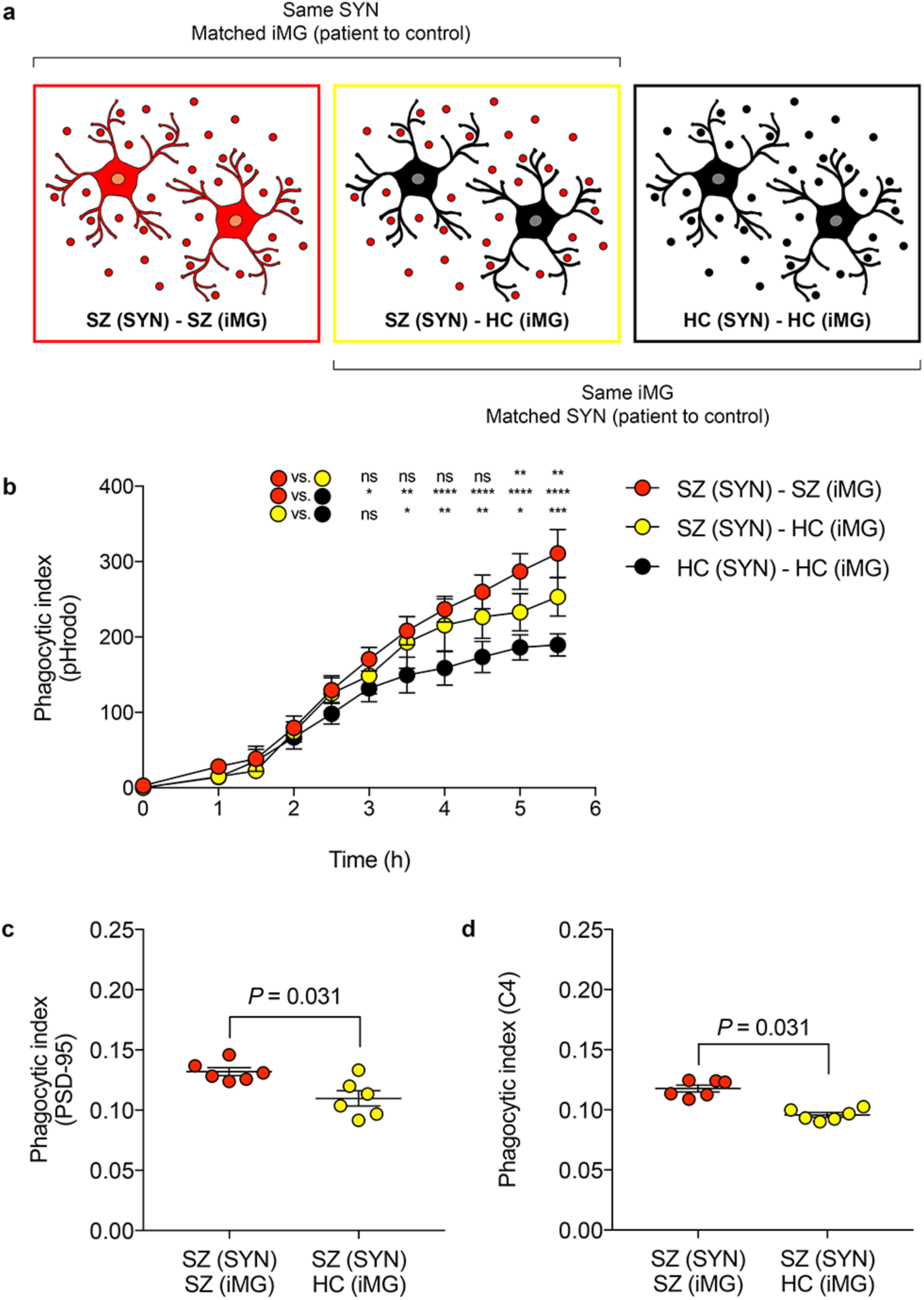
Microglial factors influence synapse engulfment. (**a**) “Pure” disease models, derived from subjects with schizophrenia (SZ), were compared to “mixed” models in which the same synaptic structures from SZ subjects were added to induced microglia-like cells (iMGs) derived from matched healthy controls (HCs), as well as to matched “pure” HC models. (**b**) Quantification of pHrodo (red)-labeled synaptosomes (SYN) uptake in iMGs during live imaging sessions of 5.5 h. SZ – SZ models (red) were based on SYN from 2 SZ patients and iMGs from 8 SZ subjects, while SZ – HC models (yellow) were based on the SYN from SZ patients and iMGs derived from 8 matched HCs. HC – HC models (black) were based on iMGs from same subjects and 3 matched neural HC lines. (**c**) Quantification of phagocytic inclusions (PSD-95^+^ inclusions, 0.5–1μm) using confocal microscopy and SYN derived from 2 neural SZ lines matched with iMGs from 6 SZ patients (red) or 6 matched HCs (yellow). All reported p-values are two-sided and error bars represent SEM. Data in (c) were analyzed using a two-way repeated repeated measures ANOVA with Sidaks’s post tests, and data in (d) were analyzed using Wilcoxon signed rank test.

### Minocycline decreases synapse engulfment *in vitro*

Minocycline, a semisynthetic brain-penetrant tetracycline antibiotic, has been suggested to be beneficial in neurodevelopmental and neurodegenerative diseases because of putative anti-inflammatory effects ^32^. While the mechanism of action is unknown, it has been postulated that minocycline works by targeting synaptic remodeling ^33^. Consistent with this hypothesis, repeated high-dose intraperitoneal administrations of minocycline in postnatal mice cause a decrease in engulfment of retinal ganglion cells (RGC) inputs; a commonly used rodent model for synaptic pruning ^17^ Clinically, however, minocycline is typically given in lower doses, yielding significantly lower brain concentrations ^34,35^, and human cellular data is lacking. Guided by previous rodent studies ^36,37^ we pre-treated iMG cultures with minocycline in a series of clinically relevant doses (1 μM, 10 μM, and 60 μM). Real-time live imaging revealed a dose-dependent decrease in engulfment of synaptosomes with nearly undetectable synaptosome uptake at 60 μM (*P* < 0.0001; **Fig. 5a**).

**Figure 5.**
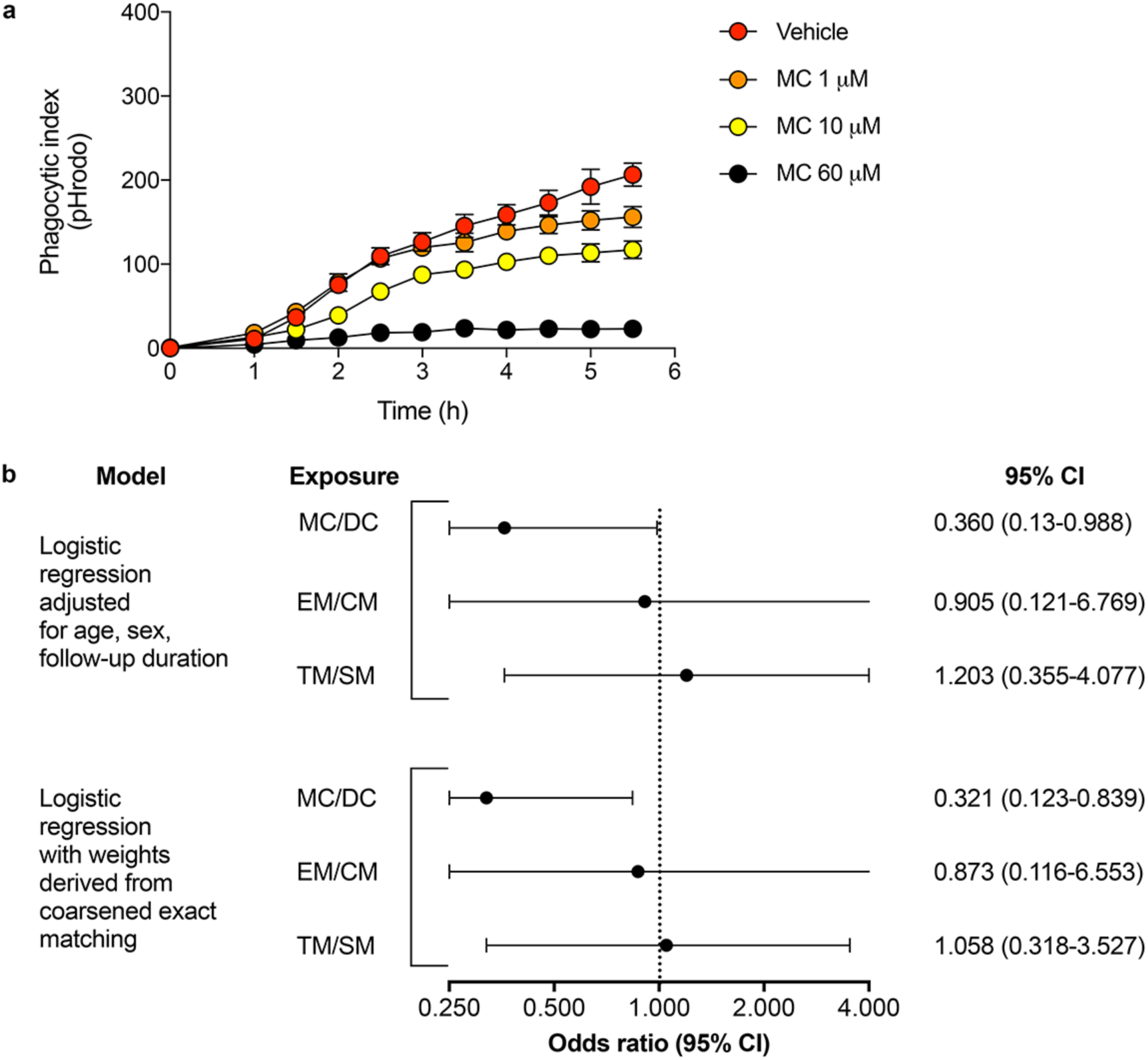
Minocycline inhibits synapse engulfment *in vitro* and decrease schizophrenia risk in electronic health records. (**a**) Quantification of pHrodo (red)-labeled synaptosomes uptake in induced microglia-like cells (iMGs) during live imaging sessions of 5.5 h. iMG cultures (n=8) were pre-treated for 30 minutes with vehicle or minocycline at a concentration of 1μM, 10μM, or 60μM. (**b**) Risk of developing schizophrenia (SZ) after exposure to minocycline or doxycycline (MC/DC), erythromycin or clindamycin (EM/CM), or trimethoprim/sulfamethoxazole (TM/SM). All data derived from an electronic health record derived longitudinal cohort of 9,031 young adults who received at least 90 days of continuous prescribed antibiotic treatment during the study period. Error bars represent SEM. All p-values two-sided. Data in (A) were analyzed using a two-way repeated measures ANOVA with Sidak’s post hoc tests.

### Drug exposure during adolescence and risk of later-on schizophrenia

As our data demonstrated that minocycline dose-dependently decreases microglial engulfment of synaptic structures in human models at clinically relevant doses, we hypothesized that chronic exposure to minocycline, or the chemically and mechanistically-similar brain-penetrant drug doxycycline, during adolescence, i.e., a time-period of intense synaptic pruning ^18^, might decrease schizophrenia risk. These two medications are among several antibiotics commonly prescribed for treating acne vulgaris ^38^, allowing a pseudo-randomized investigation using the electronic health records of a large academic medical center. Given the challenges in terms of feasibility of prospective randomized studies, we sought to identify proof-of-concept that might motivate such next-step investigation. To avoid confounding by indication, we analyzed 9,031 individuals age 18 or younger, treated for acne but without a prior diagnosis of schizophrenia or schizoaffective disorder, who received at least 90 days of continuous electronically-prescribed antibiotic use during the study period, contrasting individuals (54 % females) with minocycline or doxycycline exposure to those with no such exposure but equivalent exposure to another antibiotic, with up to 8 years of follow-up (mean follow-up time 5.6 years [SD 3.6]). Exposure to minocycline or doxycycline was associated with a significantly decreased risk for incident schizophrenia (odds ratio [OR] = 0.363, 95% CI: 0.133-0.988; **Fig. 5b**) during follow-up. For comparison, we also studied erythromycin and clindamycin, as well as trimethoprim/sulfamethoxazole, using identical designs. Unlike minocycline and doxycycline, we did not observe a decreased risk for schizophrenia (erythromycin/clindamycin: OR = 1.203, 95% CI 0.355-4.077, and trimethoprim/sulfamethoxazole: OR 0.905, 95% CI 0.121-6.769; **Fig. 5b**). To confirm the robustness of the inverse association between minocycline/doxycycline exposure and schizophrenia by reducing potential imbalance in covariates between groups, we repeated the primary analysis using coarsened exact matching (CEM). Among 8,726 matched individuals, this logistic regression analysis, with weights derived from CEM, confirmed our primary observation of a decreased risk in subjects exposed to minocycline or doxycycline (OR = 0.321, 95% CI 0.123-0.839; **Fig. 5B**). Sensitivity analyses, examining the effects of minocycline and doxycycline individually, also yielded similar point estimates (minocycline, OR = 0.425, 95% CI 0.125-1.442, doxycycline, OR = 0.342, 95% CI 0.82-1.511).

## DISCUSSION

Applying cellular reprogramming methods to create patient-specific models of synaptic pruning in individuals with schizophrenia and carefully matched HCs, we demonstrate robust support for increased elimination of isolated synapses among schizophrenia-derived iMG cultures, attributable independently to neural as well as microglia-specific effects inhibited in a dose-dependent fashion by minocycline. We further report proof-of-concept clinical data in a large cohort of antibiotic-treated young adults with acne identified in electronic health records supported a reduction in incident schizophrenia risk following chronic minocycline but not other antibiotic exposure during adolescence, after adjustment for potential confounding variables.

Our experimental data using patient-derived cells offer a potential mechanism explaining structural and functional disturbances observed in schizophrenia patients. Loss of cortical gray matter volume among schizophrenia patients has been repeatedly observed ^39^. Decreased synaptic density has been reported from postmortem material ^12–14^ and functional magnetic resonance imaging (fMRI) studies display aberrant functional brain connectivity ^9–11,39^. Studies of ‘at risk’ adolescence or early adulthood populations further reveals strong evidence for an accelerated decrease in gray matter thickness among ‘converters’ ^3–8^, suggesting that the observed structural aberrations are presented already at transition to illness and are likely to be related to disturbance in cortical maturation in the given age interval. Rodent studies demonstrate that microglia, dependent on complement signaling, play a major role in synaptic pruning ^16,17,40^, a prominent feature of human neurodevelopment during adolescence ^18^, while microglial activity, as measured by [^11^C]PBR28 positron emission tomography (PET) scans, correlate to cortical gray matter loss in subjects at ultra-high risk of psychosis ^41^.

In line with data from mice deficient in *C4* ^15^, sharing sequence similarities to both human *C4A* and *C4B*, we observed a strong correlation between a schizophrenia risk variant, linked to increased *C4A* expression, and increased synapse engulfment. By combining synapses and iMGs from subjects with different *C4* haplotypes we were also able to specifically link the effect of genetically predicted increased *C4A* over *C4B* expression (controlled for total *C4* copy numbers) to neurons. While *C4* risk variants exclusively influenced synapse elimination through a neural mechanism we also observed effects on synapse engulfment through *C4*-independent effects related to intrinsic factors in iMGs. Notably, drug targets that are specifically expressed in microglia may be more favorable in terms of adverse immune effects, and studies aiming to identify such targets are warranted. Pre-treating iMGs with the antibiotic minocycline we observed a dose-dependent decrease of synaptosome uptake. This first observation in our human model system accords with a previous animal study reporting decreased engulfment of RGC inputs among mice treated with minocycline ^16^. Engulfment of viable neural progenitor cells, a microglial function that we have previously shown can be mimicked *in vitro* by iMGs ^21^, is also inhibited by minocycline using a rodent model in which an induced status epilepticus increases numbers of progenitors in the subgranular zone of the dentate gyrus ^42^. In contrast, minocycline does not decrease phagocytosis of amyloid β fibrils in precursor protein transgenic (APP-tg) mice ^43^, although it prevent secretion of pro-inflammatory cytokines ^43^, and even rescues decreased engulfment of latex beads by adult hippocampi microglia isolated from Polyinosinic:polycytidylic acid (Poly(I:C)) mice ^44^. Taken together, this work demonstrates that minocycline influence microglial phagocytic function in a highly complex but dissectible manner where factors such as activation state and given substrate may be of crucial importance.

Our pseudo-randomized investigations using electronic health records revealed a decreased risk of incident schizophrenia after chronic exposure to minocycline but not non-tetracycline antibiotics during adolescence. This finding requires confirmation in prospective randomized trials, which may be challenging in terms of feasibility but are warranted by our results. Previous clinical studies have yielded mixed effects for minocycline in small-scale clinical investigation, but importantly these studies only examined acute efficacy in chronic schizophrenia subjects. Our results suggest that minocycline has the capacity to delay, or even prevent, the onset of schizophrenia through modulation of microglia-mediated synaptic elimination. Such an effect would be of major clinical importance if confirmed prospectively, as it would represent the first demonstrated means of preventing this chronic and disabling disorder. These studies, perhaps coupled with *in vitro* investigation using our model, represent an important next step.

Several limitations in the current study should be highlighted. First, the transcriptomic signature of human *ex vivo* microglia changes rather rapidly *in vitro* ^20^. iMGs, similar to recent iPSC-derived microglia-like cells and primary cultured microglia, do not fully recapitulate the molecular signature of human *ex vivo* microglia despite adjusting conditions to mimic the brain microenvironment. However, despite these limitations our *in vitro* assays using iMGs were still able to recapitulate mechanisms observed *in vivo* using rodent models ^21^. It is also worth noting that the current study lacks direct evidence supporting that the observed effects of minocycline *in vitro* are per se *causing* the decreased incidence of schizophrenia observed in our register-based analyses. We intend the latter as proof-of-concept, but can only demonstrate association using this design. Although the time-period of minocycline exposure overlaps with cortical maturation due to extensive synaptic pruning it is fully possible that minocycline reduces risk of schizophrenia through another as-yet unidentified mechanism.

Nonetheless, taken in aggregate, our work supports increased synapse elimination in schizophrenia, offering a mechanism linking genetic risk variants to observed structural brain changes in early disease. Specific pharmacological interventions targeting microglial elimination of synapses merit further study for their capacity to delay or prevent the onset of schizophrenia in high-risk individuals.

## ONLINE METHODS

### Study design

For patients vs. controls comparison of synaptosome uptake by iMGs, using live-imaging, we assumed a between-subject effect size of 0.5 (repeated ANOVA, α = 0.05, and correlation among repeated measurements [n=10] = 0.5). With a power of 0.9 this indicated a total sample size of 26 (13 subjects per group) ^45,46^. Using 3 iPSC-derived neural lines from schizophrenia patients and 2 iPSC-derived neural lines from HCs we created 13 schizophrenia models and 13 HCs models by combining derived synaptosomes with iMGs from 13 male schizophrenia subjects and 13 male HCs. Patient vs. controls, as well as each synaptosome – iMG pair, were matched on age. To reduce technical variance nine images per well were taken and averaged. Post-assay cocultures were then stained for PSD-95 and C4 (see below). Immunohistochemistry for 2 cocultures failed due to technical issues. Synaptosomes from 2 additional schizophrenia subjects and 2 matched HCs were then derived and subjects in the Massachusetts General Hospital (MGH) Neurobank (see ‘Subjects’) with adequate matching criteria, including donors described above, were re-contacted and asked for a new blood draw in order to derive iMGs. If more than one buffy coat was obtained, or the subject already contributed with cells, we matched additional iMG cultures with synaptosomes from another neural line to maximize numbers of biological replicates in terms schizophrenia and HC models (numbers of subjects as well as cellular models are stated in each figure legend). 20 randomly selected images were taken per well and as pre-defined, only images displaying more than 20 cells and less than 100 cells were included in further analyses (previous experiments in our lab suggests that risk of outliers increases unacceptably outside this range). Technical variance between wells for a given HCs iMG line has previously been shown to be very low ^21^. To examine consistency of within-individual microglial assays in a patient vs. control setting and using different buffy coats, for 8 individuals (4 schizophrenia subjects and 4 HCs) microglia samples were now derived from two different buffy coat samples. Correlation between pairs (R-squared) of samples was 0.92 (**Supplementary Figure 6**). Investigators performing the experiments were blinded to well conditions. For analysis of spine density in co-culture experiments (see below), fields lacking identifiable neurons or spines were excluded from analysis.

### Subjects

Study participants (18-75 years old) were recruited from clinical programs at MGH, as well as by using advertisements and study volunteer lists (MGH Neurobank). All experiments were performed with informed consent prior to participation, as approved by the Institutional Review Board of Partners HealthCare. A structured psychiatric evaluation was performed for all included subjects (17 schizophrenia subjects and 18 HCs) using the structured clinical interview with both the DSM-IV (SCID) and the Mini-International Neuropsychiatric Interview (MINI) by a single psychiatrist with at least 5 years of clinical experience ^47^. Background characteristics for participants are summarized in **Supplementary Table 6.**

### Generation of induced microglia-like cells (iMGs)

iMGs were derived from monocytes using established methods as previously described in detail ^21^, but with some modifications. Briefly, peripheral blood mononuclear cells (PBMCs) generated from whole blood were rapidly thawed at 37°C and diluted into pre-warmed RPMI-1640 medium (Sigma-Aldrich, Saint Louis, MO, USA; #R0883-500ML) with 10% heat-inactivated low-endotoxin FBS (Rockland Immunochemicals, Hamburg, Germany; #FBS-01-0100) and spun at 300 x g for 5 minutes (break off). The supernatant was aspirated completely and the pellet was resuspended in RPMI-1640 medium (Sigma-Aldrich) with 10% heat-inactivated low-endotoxin FBS and 1% penicillin⤓gstreptomycin (P/S). Cells were plated onto Geltrex (Thermo Fisher Scientific, Waltham, MA, USA; #A1048002) coated 24 well plates at a density of 1 x 10^6^ cells per well and incubated overnight at 37°C. Media was then changed to RPMI 1640 medium with Glutamax (Life Technologies, Carlsbad, CA, USA) with 1% P/S, 0.μg/ml of IL-34 (R&D systems, Minneapolis, MA, USA: #5265-IL-010) and 0.01ug/ml of GM-CSF (R&D systems, #215-GM-010/CF). After 10 days the plates were washed thoroughly to remove unbound cells and fresh media was added. Cells were harvested or used for functional assays the following day.

### Collecting dermal biopsies and establishing fibroblast cultures

Following subcutaneous injection of lidocaine (1%), biopsies were obtained from the non-dominant upper arm by a physician investigator (via 3.0 or 4.0mm punch tool). A scalpel blade was used to cut the biopsy into small pieces and then transferred to a 60mm tissue-culture treated dish (Corning, NY, USA) on which the biopsy pieces were plated dermis side down. The dish was placed in an incubator at 37°C for 15 minutes in order for the dermal sample to adhere to the dish. Media that contained Dulbecco’s modified Eagle medium (DMEM; Life Technologies; #11995-065) with 10% heat inactivated FBS (Life Technologies; #10082147) and 1% penicillin–streptomycin–glutamine (Corning; #30-009-CI) was gently added over the pieces of biopsy. The resulting fibroblasts were passaged, confirmed as mycoplasma negative (MycoAlert™ Mycoplasma Detection Kit, Lonza), frozen down, and then thawed as needed for iPSC reprogramming.

### iPSC reprogramming and neural differentiation

Human fibroblasts were reprogrammed, stabilized, and expanded under xeno-free conditions by Stemiotics, Inc. (San Diego, CA) as previously described ^27^ Briefly, iPSC colonies were obtained using mRNA reprogramming in a feeder-free culture system. Stable iPSCs were expanded in Nutristem XF media (Biological Industries, Kibbutz Beit-Haemek, Israel) and on rLaminin-521 (BioLamina, Sundbyberg, Sweden; #LN521-02) coated plates) to at least passage 3. iPSCs were then purified using magnetic-activated cell sorting (MACS) with TRA-1-60 microbeads (Miltenyi Biotec, Bergisch Gladbach, Germany) on LS columns per manufacturer’s instructions. NPC induction was then initiated by media replacement to Neurobasal™ media (Gibco; #21103049) supplemented with 1 × Neural Induction Supplement (Gibco; NIS media). After confluence, cells were passaged with Accutase (Sigma-Aldrich; #A6964-100ML) and purified using PSA-NCAM MACS sorting procedure (Miltenyi Biotec). The medium was changed to a neural expansion medium (NEM; 50% Neurobasal medium, 50% advanced DMEM/F-12 with 1 × Neural Induction Supplement, Gibco) for further expansion. When confluent, putative NPCs were further purified by double sorting using MACS columns and CD271 and CD133 MACS kits (Miltenyi Biotec). CD2717^−^/CD133^+^ NPCs were analyzed for NESTIN, SOX1, SOX2 and PAX6 expression by immunocytochemistry. NPCs were further differentiated to neurons by plating expanded cells at a seeding density of 40 000 cells per cm^2^ on polyornthine/laminin coated 6-well tissue culture plates in stochastic neural differentiation medium (70% DMEM [Gibco] 30% Ham’s F-12 (Mediatech Inc., Tewksbury, MA, USA)) supplemented with B-27 (Invitrogen) with medium replacement every 3–5 days for 6 weeks.

### Global gene expression microarray hybridization and data analyses

Cells were harvested using Accutase and pelleted by centrifugation at 300 × g. The supernatant was removed and the pellets were stored at -80°C. Total RNA was extracted using the RNeasy Mini kit (Qiagen, Hilden, Germany) as per manufacturer’s instructions and concentrations and 260nm/280nm readings were determined using a NanoDrop™ 2000 Spectrophotometer (ThermoFisher Scientific). RNA quality was further assessed on an Agilent 2100 Bioanalyzer (Agilent Technologies Inc.) and samples with RNA integrity numbers (RIN) > 6 were used for analysis. Whole-genome gene expression profiling was performed using the Clariom™ S Pico human gene expression array (Affymetrix Inc., Santa Clara, CA, USA) with 4.5 ng total RNA per each sample at the Boston Children’s Hospital IDDRC Molecular Genetics Core Facility (supported by National Institutes of Health award NIH-P30-HD 18655). A GeneChip™ Fluidics Station 450 performed all liquid handling steps and chips were scanned with the GeneChip Scanner 3000 7G (Affymetrix Inc.). Quality control (QC) was performed using the manufacturer’s software Transcriptome Analysis Console (TAC) v4.0 (Affymetrix Inc.), as per manufacturer’s instructions (PCA, labeling controls, hybridization controls, Pos vs. Neg AUC, and Signal Box Plot) with no indication of outliers. Normalization was performed according to a Robust Multi-chip Analysis (RMA) approach. Differentially expressed genes were identified using a one- or two-way ANOVA. Uncorrected as well as FDR-adjusted (Benjamini-Hochberg) p-values are reported. To assess the uncertainty in the hierarchical cluster analysis of iMGs vs. monocytes and monocyte-derived macrophages (**Supplementary Fig. 3**) we applied bootstrap resampling (10,000 iterations) via the R package pvclust. For each suggested cluster pvclust provides two types of P-values: the Bootstrap Probability value from the ordinary bootstrap resampling and the Approximately Unbiased (AU) probability value from multiscale bootstrap resampling. The multiscale bootstrap resampling method was introduced to develop an approximately unbiased test, and therefore it provides better estimations of the probability values ^34^. The standard errors of AU P-values were also plotted against AU P-values with no indication of sampling error.

### Synaptic vesicle endocytosis and synapse markers in isolated nerve terminals

With some modifications, we adopted the methods of Daniel et al. ^31^ for high-throughput quantification of SVE to validate functionality of synaptosomes isolated from iPSC-derived neuronal cultures. First, 96-well plates (Ibidi, Martinsried, Germany; #89626) were coated with PEI (Sigma-Aldrich; P3143 1:15000 diluted in water) at 37°C overnight. The next day, plates were flicked to remove excess PEI, further dried at 37°C for 30 minutes and stored at 4°C for a maximum of 30 days. Frozen synaptosomes were thawed and sonicated for one hour at room temperature. 200 μl of sucrose/EDTA/Tris buffer (SET) buffer (0.32 M sucrose, 1mM EDTA, and 5 mM Tris dissolved in water and adjusted for pH 7.4) was added to the synaptosomes and mixed thoroughly. 1:1000 DTT (Sigma, #D5545; 250 mM stock solution dissolved in SET buffer) solution was added to the synaptosome solution, and 100 μl of the final (300 μl) solution was added to each well to give triplicate data points. The plate was centrifuged at 1500 x g for 30 minutes at 4°C. HEPES-buffered Krebs-like (HBK) buffer (143 mM NaCl, 4.7 mM KCl, 1.3 mM MgSO_4_.7H_2_O, 1.2 M CaCl_2_, 0.1 mM Na_2_HPO_4_ and 10 mM D-glucose dissolved in water and adjusted at pH 7.4) was pre-warmed at 37°C and infused with 5% CO_2_ for 30 minutes. After centrifugation of synaptosomes, SET buffer was carefully removed using a multichannel pipette and 60μl of HBK buffer was added. The synaptosomes were then incubated at 37°C for 15 minutes to acclimate to HBK. Simultaneously, a solution of 1.5μM Calcein Blue-AM (Invitrogen, #C1429) was prepared in HBK buffer. After incubation, HBK buffer was carefully removed and Calcein Blue-AM (60μl per well) was added followed by incubation for 30 minutes at 37°C in the dark. FM4-64 dye (Invitrogen, #T13320) was diluted 1:100 in water for unstimulated control wells, and diluted 1:100 in 400mM KCl for stimulated wells. These solutions were protected from light. After incubation with Calcein Blue-AM, synaptosomes were removed from the incubator and the solutions were carefully replaced with 81 μl of HBK buffer. Starting with the wells for unstimulated controls, 9μl of the unstimulated solution was added to the unstimulated wells, followed by 9μl of stimulated solutions for the stimulated wells, using a multichannel pipette. The plate was incubated for two minutes at 37°C. Media was aspirated carefully with a pipette and replaced by 81 μl of HBK buffer and 9 μl of ADVASEP-7 (Sigma-Aldrich, St. Lois, MO, USA, #A3723) was added to stop the depolarization and wash off the excess FM4-64 from the plasma membranes. The plate was incubated for another two minutes at 37°C, after which ADVASEP-7 solution was replaced by 90μl of HBK using a multichannel pipette. The plate was then immediately imaged using an IN Cell 6000 confocal analyzer (GE Healthcare). 40 randomly selected images were acquired per well with all settings kept constant between conditions. After Imaging, plates were fixed with 4% PFA for 15 minutes at room temperature. The synaptosomes were washed carefully with PBS and incubated with anti-PSD-95 antibody conjugated to Alexa Flour 488 (Abcam #195004) for 20 minutes at room temperature. Wells were then washed once with PBS and imaged with on the IN Cell 6000 using the same settings (including same areas of the well) as FM4-64. Quantification was performed using the IN Cell Analyzer 1000 software (GE, Healthcare). The number of PSD-95^+^ puncta (0.5-1μm) co-localized with FM4-64 was quantified on each images of the 40 images per well and used to create a mean count (over three wells per line) representative of concentration of PSD-95^+^ synaptosomes for a specific line. To makes sure that our main analyses were not substantially influenced by selective bias due to differences in harvested synaptosome counts, we also performed all main analyses adding these counts as covariates (multiple logistic regression). Given limited line-to-line variability with a random distribution between cases and controls, all adjusted analyses displayed similar effect sizes and significances for the different phagocytic index (data not shown).

### Transmission electron microscopy (TEM)

Isolated synaptosome suspensions were fixed with 2.0% glutaraldehyde in 0.1M sodium phosphate buffer (pH 7.4, Electron Microscopy Sciences, Hatfield, PA, USA) for two hours at room temperature, then centrifuged at 14 000 rpm for 10 minutes to obtain a working pellet. Pellets were infiltrated in 1.0% osmium tetroxide in cacodylate buffer for one hour at room temperature and gently rinsed several times in cacodylate buffer. Pellets were stabilized in 2% agarose, and agarose blocks (containing pelleted material) were dehydrated through a graded series of ethanols to 100%, then dehydrated briefly in 100% propylene oxide. Specimens were pre-infiltrated for two hours in a 2:1 mix of propylene oxide and Eponate resin (Ted Pella, Redding, CA, USA), then transferred into a 1:1 mix of propylene oxide and Eponate resin and allowed to infiltrate overnight on a gentle rotator. The following day, specimens were allowed to infiltrate in fresh 100% Eponate resin for several hours, embedded in flat molds with 100% Eponate, and allowed to polymerize 24 to 48 hours at 60°C. Thin (70 nm) sections were cut using a Leica EM UC7 ultramicrotome, collected onto formvar-coated grids, stained with uranyl acetate and Reynold’s lead citrate and examined in a JEOL JEM 1011 Transmission Electron Microscope at 80 kV. Images were collected using an AMT digital imaging system (Advanced Microscopy Techniques, Danvers, MA, USA). Electron microscopy was performed in the Microscopy Core of the Center for Systems Biology/Program in Membrane Biology at MGH, which was partially supported by an Inflammatory Bowel Disease Grant DK043351 and a Boston Area Diabetes and Endocrinology Research Center (BADERC) Award DK057521.

### Co-culture system

Co-culture experiments were conducted with iPSC-derived neurons following a 90-day cortical neural differentiation protocol adapted from previously described ^48^. Briefly, iPSCs were grown in N2B27 supplemented neurobasal medium containing LDN 193189 (100 nM; Stemgent, USA) and SB 431542 (10 μM; Stemgent, USA) with daily media changes for 10 days then switched to N2B27 medium alone for the remainder of the differentiation. Neural cultures were replated into 24 well plates at a density of 2.5 x 10^4^ cells / cm^2^ in 0.5 mL N2B27 on day 80 and further cultured for 10 more days. Neurons were pHrodo labeled for one hour at 37°C followed by a media change. Mature iMGs were detached from plates using Accutase and added to neural cultures. The co-culture plates were then imaged using Incucyte ZOOM live-imaging system (Essen Biosciences) with images collected every 4 hours for a period of 48 hours. After assay, cells were washed and fixed with 4% PFA for 15 minutes and stained with Alexa Fluor™ 647 Phalloidin (Life Technologies Inc.; #A22287) as per manufacturer’s instructions. Cells were then imaged on the IN Cell 6000 confocal analyzer (GE Healthcare) as described above. Raw 16-bit TIFF files containing image data were converted to RGB images using ImageJ (NIH, MD, USA). From these images, 10μm sections of phalloidin-stained dendritic spines were identified, cropped, and opened by an operator (B.W.) blinded to well conditions in SynPAnal for spine analysis ^49^. The 10μm dendrite regions were then highlighted, and spine regions were identified and counted automatically. Lastly, the numerical data exported to Libreoffice Calc spreadsheet (The Document Foundation, Germany) for analysis. Further inferential statistics were performed using Graphpad Prism 6.0 (Graphpad Software Inc., CA, USA). 70 fields of view were analyzed per well with one process selected per field.

### Isolating and labeling nerve terminals

24 hours before harvesting neural cultures by scraping, medium was changed to ACM (ScienCell, Carlsbad, CA, USA). Synaptosomes were then isolated as previously described using Synaptic Protein Extraction Reagent (Syn-PER; Thermo Fisher Scientific) ^21^. Resulting synaptosomes were collected and labeled with an amine-reactive and pH-sensitive dye (pHrodo Red, SE; Thermo Fisher Scientific; #P36600) as per manufacturer’s instructions.

### Western blotting analyses

Synaptosomes were centrifuged at 12000 × g for 15 minutes and pellets were re-suspended in radioimmunoprecipitation assay (RIPA) buffer with 0.2% Triton containing protease inhibitors (Roche, Basel, Switzerland; #05892791001), kept on ice for 15 minutes and centrifuged again at 20000 × g for 10 minutes. The supernatant was then loaded onto a Biorad TGX 4-20% Mini-PROTEAN ™ Precast gel (Biorad, Hercules, CA, USA; #4561096) and run at 120V. The gel was then transferred onto an Immobilon-FL PVDF membrane (Millipore, Billerica, MA, USA, #IPFL00010) at 120 V for 1 hour. After transfer, the membrane was blocked for one hour with licor blocking buffer (LI-COR Biosciences, Lincoln, NE, USA, #927 40100). The blot was then cut and incubated with primary antibodies: anti-Synaptophysin (1:7500, Abcam; #ab32127), anti-SNAP-25 (1:1000, Abcam; #ab53723), anti-C4 (1:1000, Abcam; #ab173577), or Beta Actin loading control (1:1000, Abcam; #ab8226) overnight at 4°C on a shaker. The blot was washed three times with phosphate-buffered saline (PBS) containing 0.1% Tween and incubated for one hour at room temperature with secondary antibodies (IRDye 800CW anti-rabbit; LI-COR Biosciences; #NC0809364) diluted 1:10000 in a buffer solution containing 50% LI-COR blocking buffer, 50% PBS, 0.1% Tween, and 0.02% SDS). After incubation, blots were washed three times with PBS containing 0.1% Tween and imaged using a LI-COR Odyssey CLx system to confirm that the synaptosomes were positive for typical synaptic markers.

### Real-time live imaging

We used a live cell imaging system, Incucyte ZOOM system (Essen Biosciences) incubated at 37°C and 5% CO_2_, as previously described ^21^, but with some modifications regarding image analyses. Briefly, sonicated and pHrodo labeled synaptosomes were added to iMG cultures on 24-well plates and imaged at a resolution of 0.61μm/pixel. Images (1392×1040 pixels) were further pre-processed in IncuCyte ZOOM software (2016A) and exported as 16 bit grayscale PNG files. For minocycline experiments, iMGs were pretreated with μM, 10μM, 60μM, or vehicle for 30 minutes before initiation of the phagocytosis assay. Data was further analyzed in MATLAB (MathWorks, R2014b) to quantify phagocytized particles. Briefly, unsharp masking using a 35 pixels standard deviation Gaussian filter was used to remove background from the images, followed by distribution-based thresholding. For each image, the threshold was calculated as 10 standard deviations above the average value for the local maxima. A threshold was applied to the resulting segmented regions to remove particles smaller than 10 pixels in area. The area of the remaining particles was summed to produce the total amount of phagocytized particles. This sum was divided on number of iMGs per image and represents phagocytic index.

### Immunocytochemistry

Cells were fixed in 4% paraformaldehyde (PFA) for 15 minutes at 4°C. After fixation, cells were washed two times with PBS and then blocked for one hour with a solution containing 5% FBS and 0.3% triton in PBS. Cells were then washed three times for 5 minutes each with 1% FBS in PBS and incubated with primary antibodies (details in **Supplementary Table 7**) for two hours at room temperature. After incubation, the cells were washed three times for 5 minutes each with 1% FBS PBS and incubated for one hour at room temperature in mixture of appropriate secondary antibodies (1:500) and Hoechst (1:5000) diluted in 5% FBS in PBS. Cells were then washed again three times for 5 minutes each with 1% FBS in PBS and imaged. Antibody specificity was confirmed by performing ‘secondary only’ experiments (data not shown).

### Confocal microscopy

Images were acquired using an IN Cell 6000 analyzer (GE Healthcare). 20 randomly selected images were acquired per well (24-wells plates) with all settings kept constant between different conditions. Quantification was performed using the IN Cell Analyzer 1000 (GE Healthcare). Nuclei were identified using a Top-Hat transform, while cells were identified using Region Growing segmentation. PSD-95, pHrodo, or C4^+^ particles were identified using a multiscale Top-Hat transform (10 transformations), a size filter of 0.5-1μm, minimum sensitivity for detection of inclusions, and restriction of detection of inclusions to cytoplasm area. For outcome measurements we counted number of identified particles per iMG cell and divided on cytoplasm area to create a phagocytic index. The mean of phagocytic indexes for all images per well was then used in further analyses.

### Molecular analyses of *C4* alleles

Similar to Sekar et al. ^15^, we measured copy numbers of each individual *C4* structural allele (*C4A*, *C4B, C4L*, and *C4S*) using ddPCR, on genomic DNA isolated from blood samples from a total of 130 controls and 54 patients with schizophrenia from the MGH Neuropsychiatric Biobank. In summary, 50ng of genomic DNA was fragmented by digestion with the restriction enzyme *AluI* ^15^. Copy numbers of *C4A, C4B, C4L and C4S* were then quantified from genomic DNA using digital droplet PCR on a QX200™ Droplet Digital™ PCR System (Bio-Rad) using specific and reference primers and probes as described by Wu et al. ^50^. Data was analyzed using QuantaSoft software (Bio-Rad), which estimated absolute copies of *C4* allele-specific DNA templates by comparison to the reference *RPP30* template present as two copies per genome.

### Risk of schizophrenia in electronic medical records

We utilized de-identified data from the electronic health records of the Partners HealthCare system, which spans multiple academic medical centers in eastern Massachusetts. Reliability of psychiatric phenotypes and medication exposures has previously been demonstrated for genetic and longitudinal outcome studies ^51–54^, including pharmacovigilance studies of similar design ^55–57^. Medication data reflected electronic prescribing by any clinician, and included duration of prescription and number of refills. We have previously demonstrated that electronic prescribing is strongly associated with presence of a detectable blood level ^58^. Of note, in the present analysis, diagnostic misclassification and medication non-adherence would both be expected to bias results toward the null hypothesis, i.e., no effect of treatment with minocycline on schizophrenia risk. The at-risk cohort was defined as individuals younger than age 18 at index visit without an ICD-9-CM diagnosis of schizophrenia (295.x) or schizoaffective disorder (295.7) who received at least 90 days of continuous antibiotic treatment for acne vulgaris. Follow-up continued until diagnosis of schizophrenia or schizoaffective disorder, or December 31, 2016, whichever came first. A priori, we required a minimum of 90 days exposure to a given antibiotic, for three reasons. First, this exposure requires documentation of at least two prescriptions, increasing confidence in adherence. Second, very short durations could still reflect other indications for antibiotic use, while treatment for this duration is more typical in acne patients ^59^. Finally, the inclusion of only antibiotic-treated patients minimizes risk of confounding by indication (for example, if another chronic infection were associated with schizophrenia risk, or non-antibiotics were preferred in a particular patient subgroup). All analyses were performed using logistic regression, to allow adjustment for potential confounding features including age and sex. Regression with coarsened exact matching ^60^ was also applied to confirm robustness of association to potential sources of bias by better balancing exposed and unexposed groups.

### Statistics

All analyses were performed using R statistics (R Development Core Team, Vienna, Austria), Graph-Pad prism 6.0 (GraphPad Software, Inc.), with the EHR data extracted and transformed using i2b2 software (i2b2 v1.6, Boston, MA, USA). Affymetrix data QC was performed using the Transcriptome Analysis Console (TAC) Software (v4.0; Thermo Fisher Scientific) as described above. Graphs were produced in Graph-Pad prism 6.0, R Statistics, or in TAC. The assumptions of each test were checked. Type of statistical test is reported in main text or in figure legends. All reported p-values are two sided.

### Data Availability

Gene expression data will be available for download from the NCBI Gene Expression Omnibus at SuperSeries XXX. Additional data that support the findings of this study are available from the corresponding authors upon reasonable request.

## ACKNOWLEDGMENTS

We thank the study participants. We are also grateful to Jayla Ruiliera (MGH) for technical assistance with isolating buffy coats. This work was supported by P50-MH106933 (NIMH and NHGRI) to Dr. Perlis, R01AT009144 (NCCIH) to Dr. Perlis and Knut och Alice Wallenbergs Stiftelse to Dr. Sellgren.

## AUTHOR CONTRIBUTIONS

C.S.M, S.D.S, and R.H.P. conceived the research. C.S.M., S.D.S., and R.H.P. contributed to the overall design, direction and reporting of the study. T.F. derived neural cultures. C.S.M. and J.G., with help from J.M.T., generated induced microglia, isolated synaptosomes, and performed all experiments as well as data analyses if not otherwise specified. B.W and R.K performed coculture experiments. J.W. and C.S.M performed data analyses of images required using Essen software. J.G. and J.M.T. performed western blotting experiments. C.P.G. performed molecular analyses of *C4* alleles. S.D.S. provided cell reprogramming expertise and R.H.P. performed analyses using electronic health records. All authors discussed the results and implications and commented on the manuscript at various stages.

## COMPETING FINANCIAL INTERESTS

The authors declare no competing financial interest.

## REFERENCES

1. Kahn, R.S., et al. Schizophrenia. Nat Rev Dis Primers 1, 15067 (2015).

2. Schizophrenia Working Group of the Psychiatric Genomics, C. Biological insights from 108 schizophrenia-associated genetic loci. Nature 511, 421–427 (2014).

3. Cannon, T.D., et al. Progressive reduction in cortical thickness as psychosis develops: a multisite longitudinal neuroimaging study of youth at elevated clinical risk. Biol Psychiatry 77, 147–157 (2015).

4. Pantelis, C., et al. Neuroanatomical abnormalities before and after onset of psychosis: a cross-sectional and longitudinal MRI comparison. Lancet 361, 281–288 (2003).

5. Ziermans, T.B., et al. Progressive structural brain changes during development of psychosis. Schizophr Bull 38, 519–530 (2012).

6. Sun, D., et al. Progressive brain structural changes mapped as psychosis develops in ‘at risk’ individuals. Schizophr Res 108, 85–92 (2009).

7. Borgwardt, S.J., et al. Reductions in frontal, temporal and parietal volume associated with the onset of psychosis. Schizophr Res 106, 108–114 (2008).

8. Takahashi, T., et al. Progressive gray matter reduction of the superior temporal gyrus during transition to psychosis. Arch Gen Psychiatry 66, 366–376 (2009).

9. Meyer-Lindenberg, A.S., et al. Regionally specific disturbance of dorsolateral prefrontal-hippocampal functional connectivity in schizophrenia. Arch Gen Psychiatry 62, 379–386 (2005).

10. Lawrie, S.M., et al. Reduced frontotemporal functional connectivity in schizophrenia associated with auditory hallucinations. Biol Psychiatry 51, 1008–1011 (2002).

11. Stephan, K.E., Friston, K.J. & Frith, C.D. Dysconnection in schizophrenia: from abnormal synaptic plasticity to failures of self-monitoring. Schizophr Bull 35, 509–527 (2009).

12. Glausier, J.R. & Lewis, D.A. Dendritic spine pathology in schizophrenia. Neuroscience 251, 90–107 (2013).

13. Glantz, L.A. & Lewis, D.A. Decreased dendritic spine density on prefrontal cortical pyramidal neurons in schizophrenia. Arch Gen Psychiatry 57, 65–73 (2000).

14. Konopaske, G.T., Lange, N., Coyle, J.T. & Benes, F.M. Prefrontal cortical dendritic spine pathology in schizophrenia and bipolar disorder. JAMA Psychiatry 71, 1323–1331 (2014).

15. Sekar, A., et al. Schizophrenia risk from complex variation of complement component 4. Nature 530, 177–183 (2016).

16. Stevens, B., et al. The classical complement cascade mediates CNS synapse elimination. Cell 131, 1164–1178 (2007).

17. Schafer, D.P., et al. Microglia sculpt postnatal neural circuits in an activity and complement-dependent manner. Neuron 74, 691–705 (2012).

18. Petanjek, Z., et al. Extraordinary neoteny of synaptic spines in the human prefrontal cortex. Proc Natl Acad Sci U S A 108, 13281–13286 (2011).

19. Cannon, T.D. How Schizophrenia Develops: Cognitive and Brain Mechanisms Underlying Onset of Psychosis. Trends Cogn Sci 19, 744–756 (2015).

20. Gosselin, D., et al. An environment-dependent transcriptional network specifies human microglia identity. Science 356(2017).

21. Sellgren, C.M., et al. Patient-specific models of microglia-mediated engulfment of synapses and neural progenitors. Mol Psychiatry 22, 170–177 (2017).

22. Muffat, J., et al. Efficient derivation of microglia-like cells from human pluripotent stem cells. Nat Med 22, 1358–1367 (2016).

23. Pandya, H., et al. Differentiation of human and murine induced pluripotent stem cells to microglia-like cells. Nat Neurosci 20, 753–759 (2017).

24. Abud, E.M., et al. iPSC-Derived Human Microglia-like Cells to Study Neurological Diseases. Neuron 94, 278–293 e279 (2017).

25. Takata, K., et al. Induced-Pluripotent-Stem-Cell-Derived Primitive Macrophages Provide a Platform for Modeling Tissue-Resident Macrophage Differentiation and Function. Immunity 47, 183–198 e186 (2017).

26. Bennett, M.L., et al. New tools for studying microglia in the mouse and human CNS. Proc Natl Acad Sci U S A 113, E1738–1746 (2016).

27. Warren, L, et al. Highly efficient reprogramming to pluripotency and directed differentiation of human cells with synthetic modified mRNA. Cell Stem Cell 7, 618–630 (2010).

28. Li, W, et al. Rapid induction and long-term self-renewal of primitive neural precursors from human embryonic stem cells by small molecule inhibitors. Proc Natl Acad Sci U S A 108, 8299–8304 (2011).

29. Sheridan, S.D., et al. Epigenetic characterization of the FMR1 gene and aberrant neurodevelopment in human induced pluripotent stem cell models of fragile X syndrome. PLoS One 6, e26203 (2011).

30. Dhara, S.K., et al. Human neural progenitor cells derived from embryonic stem cells in feeder-free cultures. Differentiation 76, 454–464 (2008).

31. Daniel, J.A., Malladi, C.S., Kettle, E., McCluskey, A. & Robinson, P.J. Analysis of synaptic vesicle endocytosis in synaptosomes by high-content screening. Nat Protoc 7, 1439–1455 (2012).

32. Solmi, M., et al. Systematic review and meta-analysis of the efficacy and safety of minocycline in schizophrenia. CNS Spectr, 1–12 (2017).

33. Inta, D., Lang, U.E., Borgwardt, S., Meyer-Lindenberg, A. & Gass, P. Microglia Activation and Schizophrenia: Lessons From the Effects of Minocycline on Postnatal Neurogenesis, Neuronal Survival and Synaptic Pruning. Schizophr Bull (2016).

34. Hersch, S., Fink, K., Vonsattel, J.P. & Friedlander, R.M. Minocycline is protective in a mouse model of Huntington’s disease. Ann Neurol 54, 841; author reply 842-843 (2003).

35. Fagan, S.C., et al. Optimal delivery of minocycline to the brain: implication for human studies of acute neuroprotection. Exp Neurol 186, 248–251 (2004).

36. Raghavendra, V., Tanga, F. & DeLeo, J.A. Inhibition of microglial activation attenuates the development but not existing hypersensitivity in a rat model of neuropathy. J Pharmacol Exp Ther 306, 624–630 (2003).

37. Nutile-McMenemy, N., Elfenbein, A. & Deleo, J.A. Minocycline decreases in vitro microglial motility, beta1-integrin, and Kv1.3 channel expression. J Neurochem 103, 2035–2046 (2007).

38. Del Rosso, J.Q. Oral Doxycycline in the Management of Acne Vulgaris: Current Perspectives on Clinical Use and Recent Findings with a New Double-scored Small Tablet Formulation. J Clin Aesthet Dermatol 8, 19–26 (2015).

39. Birur, B., Kraguljac, N.V., Shelton, R.C. & Lahti, A.C. Brain structure, function, and neurochemistry in schizophrenia and bipolar disorder-a systematic review of the magnetic resonance neuroimaging literature. NPJSchizophr 3, 15 (2017).

40. Paolicelli, R.C., et al. Synaptic pruning by microglia is necessary for normal brain development. Science 333, 1456–1458 (2011).

41. Selvaraj, S., et al. Brain TSPO imaging and gray matter volume in schizophrenia patients and in people at ultra high risk of psychosis: An [11C]PBR28 study. Schizophr Res (2017).

42. Luo, C., Koyama, R. & Ikegaya, Y. Microglia engulf viable newborn cells in the epileptic dentate gyrus. Glia 64, 1508–1517 (2016).

43. Familian, A., Eikelenboom, P. & Veerhuis, R. Minocycline does not affect amyloid beta phagocytosis by human microglial cells. Neurosci Lett 416, 87–91 (2007).

44. Giovanoli, S., et al. Preventive effects of minocycline in a neurodevelopmental two-hit model with relevance to schizophrenia. TranslPsychiatry 6, e772 (2016).

45. Faul, F., Erdfelder, E., Lang, A.G. & Buchner, A. G*Power 3: a flexible statistical power analysis program for the social, behavioral, and biomedical sciences. Behav Res Methods 39, 175–191 (2007).

46. Faul, F., Erdfelder, E., Buchner, A. & Lang, A.G. Statistical power analyses using G*Power 3.1: tests for correlation and regression analyses. Behav Res Methods 41, 11491160 (2009).

47. Sheehan, D.V., et al. The Mini-International Neuropsychiatric Interview (M.I.N.I.): the development and validation of a structured diagnostic psychiatric interview for DSM-IV and ICD-10. JClin Psychiatry 59 Suppl 20, 22–33;quiz 34-57 (1998).

48. Shi, Y., Kirwan, P. & Livesey, F.J. Directed differentiation of human pluripotent stem cells to cerebral cortex neurons and neural networks. Nat Protoc 7, 1836–1846 (2012).

49. Danielson, E. & Lee, S.H. SynPAnal: software for rapid quantification of the density and intensity of protein puncta from fluorescence microscopy images of neurons. PLoS One 9, e115298 (2014).

50. Wu, Y.L., et al. Sensitive and specific real-time polymerase chain reaction assays to accurately determine copy number variations (CNVs) of human complement C4A, C4B, C4-long, C4-short, and RCCX modules: elucidation of C4 CNVs in 50 consanguineous subjects with defined HLA genotypes. J Immunol 179, 3012–3025 (2007).

51. Castro, V.M., et al. Validation of electronic health record phenotyping of bipolar disorder cases and controls. Am J Psychiatry 172, 363–372 (2015).

52. Perlis, R.H., et al. Using electronic medical records to enable large-scale studies in psychiatry: treatment resistant depression as a model. Psychol Med 42, 41–50 (2012).

53. Gallagher, P.J., et al. Antidepressant response in patients with major depression exposed to NSAIDs: a pharmacovigilance study. Am J Psychiatry 169, 1065–1072 (2012).

54. Gainer, V.S., et al. The Biobank Portal for Partners Personalized Medicine: A Query Tool for Working with Consented Biobank Samples, Genotypes, and Phenotypes Using i2b2. J Pers Med 6(2016).

55. Castro, V.M., et al. Stratifying Risk for Renal Insufficiency Among Lithium-Treated Patients: An Electronic Health Record Study. Neuropsychopharmacology 41, 1138–1143 (2016).

56. Castro, V.M., et al. Absence of evidence for increase in risk for autism or attention-deficit hyperactivity disorder following antidepressant exposure during pregnancy: a replication study. TranslPsychiatry 6, e708 (2016).

57. Balekian, D.S., et al. Pre-birth cohort study of atopic dermatitis and severe bronchiolitis during infancy. Pediatr Allergy Immunol 27, 413–418 (2016).

58. Roberson, A.M., Castro, V.M., Cagan, A. & Perlis, R.H. Antidepressant nonadherence in routine clinical settings determined from discarded blood samples. J Clin Psychiatry 77, 359–362 (2016).

59. Bienenfeld, A., Nagler, A.R. & Orlow, S.J. Oral Antibacterial Therapy for Acne Vulgaris: An Evidence-Based Review. Am J Clin Dermatol (2017).

60. Iacus, S.M., King, G. & Porro, G. Multivariate Matching Methods That are Monotonic Imbalance Bounding. Journal of the American Statistical Association 106, 345–361 (2011).

